# Y chromosomal noncoding RNAs regulate autosomal gene expression via piRNAs in mouse testis

**DOI:** 10.1101/285429

**Authors:** Hemakumar M. Reddy, Rupa Bhattacharya, Zeenath Jehan, Kankadeb Mishra, Pranatharthi Annapurna, Shrish Tiwari, Nissankararao Mary Praveena, Jomini Liza Alex, Vishnu M Dhople, Lalji Singh, Mahadevan Sivaramakrishnan, Anurag Chaturvedi, Nandini Rangaraj, Thomas Michael Shiju, Badanapuram Sridevi, Sachin Kumar, Ram Reddy Dereddi, Sunayana M Rayabandla, Rachel A. Jesudasan

## Abstract

Majority of the genes expressed during spermatogenesis are autosomal. Mice with different deletions of Yq show sub-fertility, sterility and sperm abnormalities. The connection between Yq deletion and autosomal gene regulation is not well understood. We describe a novel mouse Yq-derived long noncoding RNA, *Pirmy*, which shows unprecedented number of splice variants in testis. Further, *Pirmy* transcript variants act as templates for several piRNAs. We identified ten differentially expressed autosome-encoded sperm proteins in mutant mice. *Pirmy* transcript variants have homology to 5’/3’UTRs of these deregulated autosomal genes. Thus, subfertility in Y-deleted mice appears to be a polygenic phenomenon that is partially regulated epistatically by the Y-chromosome. Our study provides novel insights into possible role of MSY-derived ncRNAs in male fertility and reproduction. Finally, sperm phenotypes from the Y-deleted mice seem to be similar to that reported in inter-specific male-sterile hybrids. Taken together, this study provides novel insights into possible role of Y-derived ncRNAs in male sterility and speciation.

## INTRODUCTION

Y chromosome has come a long way from a single-gene male determining chromosome to one that houses a few protein-coding genes besides sequences crucial for spermatogenesis and fertility (Bellott et al., 2014; Jehan et al., 2007; Kuroda-Kawaguchi et al., 2001; Moretti et al., 2017; Piergentili, 2010; Soh et al., 2014; Tiepolo & Zuffardi, 1976). Earlier studies have shown that genes involved in sex determination and spermatogenesis are present on the short arm. Several lines of evidence indicate that the male-specific region on the long arm of the Y chromosome (MSYq) in mouse is replete with highly repetitive mouse-specific sequences that are expressed in spermatids (Burgoyne & Mitchell, 2007; Conway et al., 1994; Prado, Lee, Zahed, Vekemans, & Nishioka, 1992; Toure et al., 2005).

Previously published data have described two different strains of mice with partial deletions of the long arm of the Y chromosome (Yq) (Conway et al., 1994; Józefa Styrna, Bili, & Krzanowska, 2002). Mice from both the genetic backgrounds exhibit male sterile phenotypes such as subfertility, sex ratio skewed towards females, reduced number of motile sperms, aberrant sperm motility and sperm head morphological abnormalities (Conway et al., 1994). Mice with partial deletions of Yq show sperm abnormalities with less severe phenotype whereas mice with total deletion of the Yq have extensive sperm morphological aberrations and are sterile (Toure et al., 2004). This suggested the presence of multicopy spermiogenesis gene(s) on mouse Yq (Burgoyne, Mahadevaiah, Sutcliffe, & Palmer, 1992; Conway et al., 1994). Subsequently multicopy transcripts such as Y353/B, spermiogenesis-specific transcript on the Y *(Ssty)* and Sycp3-like, Y-linked *(Sly)* from the mouse Yq were projected as putative candidate genes for male sterility and spermiogenesis in mice (Cocquet et al., 2009; Conway et al., 1994; Riel et al., 2013; Toure et al., 2004). As mice with partial deletions of Yq (XY^RIII^qdel, 2/3^rd^ interstitial deletion of Yq) show reduced expression of *Ssty* and impaired fertility, this gene (present on Yq) was implicated in spermatogenesis (Toure et al., 2004). The next major gene to be discovered on mouse Yq was the multicopy *Sly*. As SLY interacts with a histone acetyl transferase and is an acrosomal protein, the authors suggested that *Sly* could control transcription and acrosome functions (Reynard, Cocquet, & Burgoyne, 2009). Further, Cocquet and colleagues observed major problems in sperm differentiation when they disrupted functions of the *Sly* gene by transgenic delivery of siRNA to the gene. Therefore, *Sly* was conjectured as a putative candidate gene for spermiogenesis (Reynard et al., 2009). However, subsequently Ward and colleagues showed that *Sly* expression alone is not sufficient for spermiogenesis (Riel et al., 2019).

Vast majority of the genes required for spermatogenesis and spermiogenesis are non-Y-linked (Choi et al., 2007; Schultz, Hamra, & Garbers, 2003; Xiao, Tang, Yu, Gui, & Cai, 2008). Deletions of the Y-chromosome leading to different degrees of male infertility prompted us to hypothesise interactions between Y-derived transcripts and autosomal genes in male fertility. Earlier, studies in the lab reported a testis-specific chimeric transcript, generated by trans-splicing between the *CDC2L2* mRNA from chromosome number 1 and a long noncoding RNA from the Y chromosome in human (Jehan et al., 2007). We hypothesized more such interactions between protein coding genes on autosomes and noncoding RNAs from the Y chromosome. In this context, we studied a mutant mouse, which had a partial deletion of the Y-chromosome heterochromatic block XY^RIII^qdel (Conway et al., 1994) to look for regulatory elements, if any, in the deleted region.

Previous studies in the lab identified 300-400 copies of a mouse Y chromosome specific genomic clone, M34 (DQ907163.1) using slot blot, sequencing and bioinformatics analyses (Bajpai, Sridhar, Reddy, & Jesudasan, 2007; Singh, Panicker, Nagaraj, & Majumdar, 1994). There is a reduction of M34 copies in the XY^RIII^qdel mice that exhibit multiple sperm abnormalities. As deletions of the Y chromosome show sperm abnormalities, we reasoned that these repeat sequences could have important functional role(s) in the multistep developmental process of sperm production. The XY^RIII^qdel mice also showed reduced transcription of M34 compared to its wild type XY^RIII^ mice in testis. In order to understand putative functions of this sequence, first of all we identified a transcript corresponding to M34, *Pirmy*, from mouse testis. Subsequent experiments identified multiple splice variants of this transcript. Parallel experiments identified deregulated proteins in the sperm proteome of the XY^RIII^qdel mice. Interestingly, genes corresponding to all these proteins localized to different autosomes. Further, we showed that the UTRs of these genes bear homology to piRNAs derived from *Pirmy*. Thus, our results demonstrate for the first time (i) a novel noncoding RNA *(Pirmy)* on mouse Y long arm (ii) large number of splice variants of *Pirmy*, and the generation of piRNAs from it in mouse testis and (iii) their putative role in regulation of autosomal genes involved in male fertility and reproduction.

## RESULTS

### M34 is transcribed in mouse testis

To address the precise function of the M34 transcript we confirmed the localization of the sex and species-specific repeat M34 on mouse Y long arm again by fluorescence in situ hybridization (FISH) (Fig. 1*A*). BLAST analysis of M34 sequence against mouse whole genome also showed maximum similarity to Y chromosome (>97% identity) with few hits on the X (Fig. 1*B*, NCBI build 38.1). M34 was then analysed for expression in adult mouse testis. FISH using M34 as a probe revealed abundant transcription in testis (Fig. 1*C*). Pretreatment with RNase abolished these signals confirming the presence of RNA (Fig. 1*D*). Expression profiling by FISH in testes showed the presence of M34 transcripts in 18-day embryos, newborns and 1-month old mice (30 days postpartum) (Fig. S1), suggesting transcription of this repeat in mouse testis from early developmental stages.

**Figure 1.**
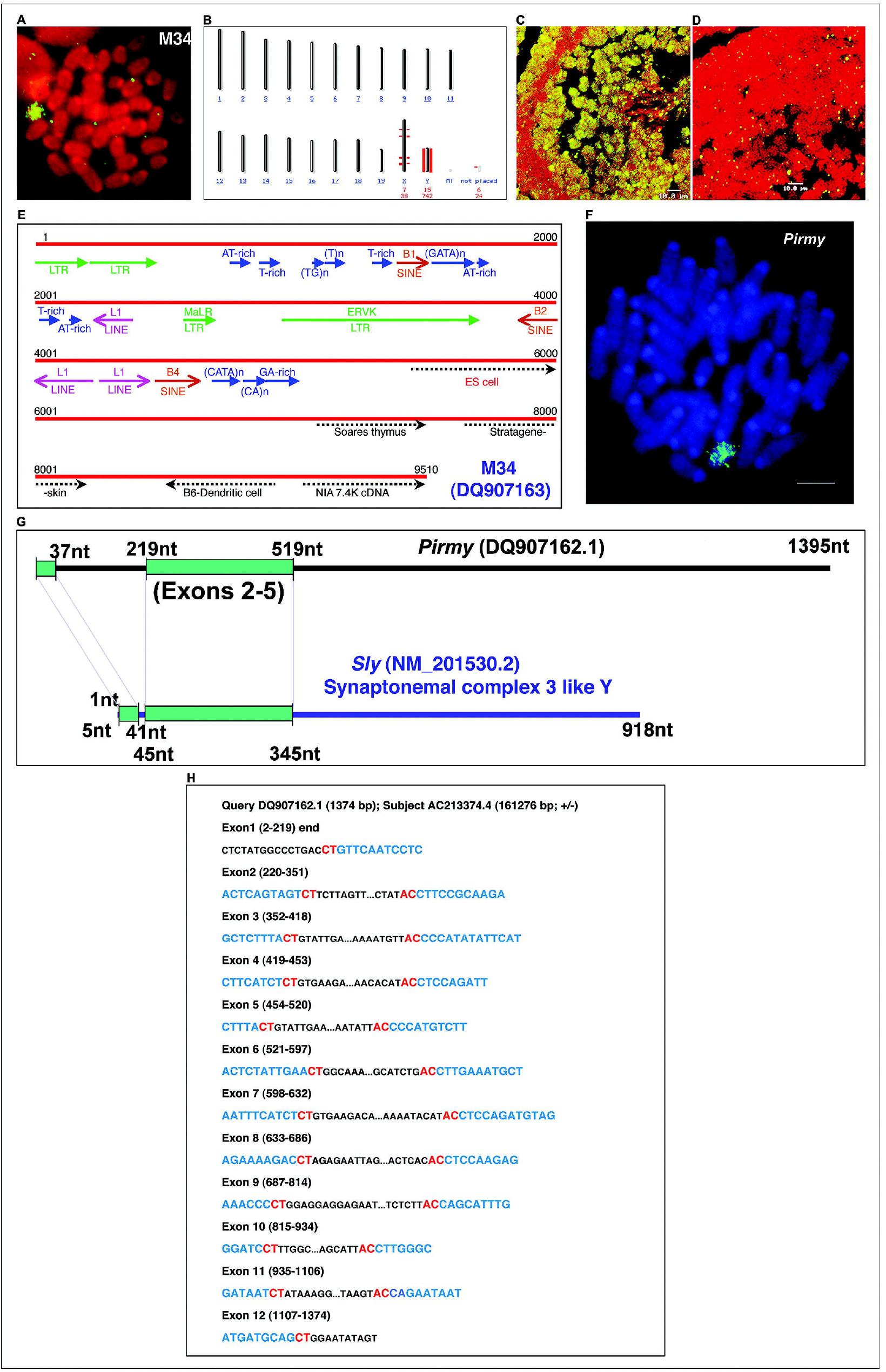
Analysis of M34 (DQ907163) and identification of a novel noncoding RNA. (*A*) Localization by FISH of the genomic clone, M34 to Mouse Y long arm in multiple copies spanning its entire length. (*B*) Mouse genome map view of M34 BLAST hits, showing Y-chromosomal localization further indicating male-specificity of these repeats. (*C*) Fluorescent in situ hybridization (FISH) using M34 shows intense signals in adult mouse testis. (*D*) Hybridization of M34 onto RNase treated testis sections does not elicit signals, indicating that the signals in panel C are due to the presence of M34 derived RNA (*E*) Sequence analysis of the 9.5kb M34 shows presence of incomplete copies of different repeats like long terminal repeats (LTRs), long interspersed nuclear elements (LINEs), short interspersed nuclear elements (SINEs), endogenous retroviral sequences (ERVK) and simple sequence repeats in both direct and reverse orientations in the clone. ESTs matching to M34 are marked as dotted arrows at the 3’end. (*F*) Shows the Y-specific localization of *Pirmy* cDNA on a mouse metaphase spread by FISH. (*G*) Partial homology between *Sly* and *Pirmy* (DQ907162) indicating identification of a novel cDNA. Homology region is highlighted in green rectangles. (*H*) Shows the consensus splice signal sequences at all intron-exon junctions. Since DQ907162 localizes to the complementary strand, the splice site consensus is seen as CT/AC instead AG/GT (see also Data Sheet).

### Isolation of a novel polyadenylated noncoding transcript from mouse Y chromosome

Sequence analysis of the 9.51 kb M34 clone using Tandem Repeats Finder (TRF) and RepeatMasker identified different simple sequence repeats and partial mid-repetitive sequences like LINEs, SINEs and LTR elements which constitute ∼35% of the total M34 sequence (Fig. 1*E*). A number of gene prediction programs like GENSCAN, GrailEXP, MZEF, and GeneMark did not predict any genes within M34 with consistency. BLAST analysis of M34 sequence against the EST database of NCBI (NCBI build 36.1) identified 5 ESTs at the 3’ end of M34 sequence (Fig. 1*E*). Expression of these ESTs was observed in embryos from at least 13.5d onwards (data not shown). Two of these ESTs showed male-specific expression by Reverse transcription PCR (RT-PCR) analysis.

In order to identify a cDNA corresponding to M34 in testis, one of the male specific ESTs (CA533654) was used to screen a mouse testis cDNA library. A 1395 nt long polyadenylated Y-specific cDNA was isolated and this was named *Pirmy* - piRNA from mouse Y chromosome (DQ907162.1). FISH on to mouse metaphase spreads showed that *Pirmy* is present only on the Y chromosome in multiple copies (Fig. 1*F*), similar to that of the genomic clone M34. BLAST of *Pirmy* against the nucleotide database of NCBI picked up only mouse sequences with statistically significant alignments (e-value <4e^-04^). This suggests that *Pirmy* is specific to mouse. BLAST analysis using *Pirmy* against the mouse genome plus transcriptome database showed homology to *Sly* transcript in exons 1-5. The exons 2-5 were identical in *Pirmy* and *Sly* (Fig. 1*G*). The region of homology is less than a quarter of *Pirmy*, and is confined to the 5’end. The entire sequence of *Pirmy* localizes to the Y chromosome at 4341127-4381724 (GRCm39). BLAST search against the nucleotide database identified a reference sequence NR164186, which was annotated as a long noncoding RNA using evidence data for transcript exon combination from *Pirmy*. The exon-intron junctions of *Pirmy* are shown in Figure 1H.

### Splice variants of *Pirmy* in mouse testis

*Pirmy* was analyzed further by RT-PCR. Two rounds of amplification using primers to the two ends of *Pirmy* yielded multiple products in testis and brain (Fig. 2*B*). PCR products were cloned. Sanger sequencing of more than 1000 clones from testis yielded 107 transcripts (NCBI accession numbers <u>FJ541075-FJ541181),</u> besides the one obtained by screening testis-cDNA library (Figs. 2A, 3). BLAST analysis of these transcripts against the NCBI genome database (Build 39) showed that 28 of these transcripts (FJ541075-FJ541102; Fig. 3) localise to multiple regions on the mouse Y chromosome. The remaining 79 transcripts (FJ541103-FJ541181) and *Pirmy* were present at a single locus on the Y chromosome at 4341127-4381724 (GRCm39) (Fig. 2). Thus, the intron-exon organization, found in *Pirmy* and the 79 splice variants is present only at a single locus on the Y chromosome. These 79 transcripts could therefore be alternatively spliced isoforms of *Pirmy*. Each exon from *Pirmy* is present in multiple copies on the Y chromosome. The 28 transcript variants reveal that different combinations of these exons and introns are there at multiple loci on the Y. The splice variants of *Pirmy* exhibited the full spectrum of splicing patterns like exon skipping, alternative 5’ and 3’ splice sites, mutually exclusive alternative exons, intron retention and combination of different splicing events (Fig. S2). The splice variants of *Pirmy* contained consensus splice signal sequences at all the intron-exon junctions (Data sheet).

**Figure 2.**
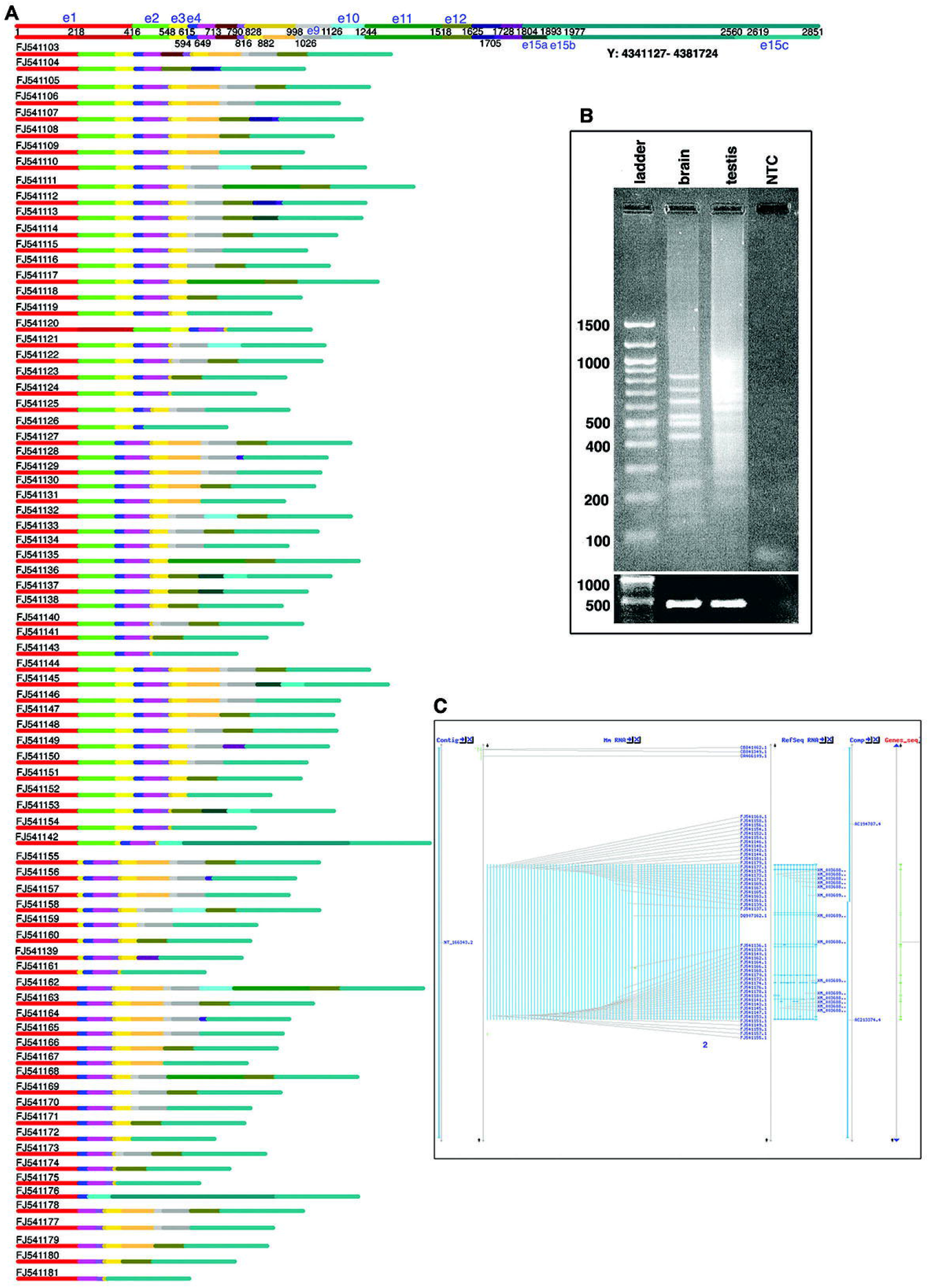
Identification of multiple splice-variants of *Pirmy*. Autoassembler program of ABI-Prism identified cDNAs that differed from one another. (*A*) Color coded line diagram showing extensive alternative splicing of *Pirmy*. The splice variants depicted here localise to Y: 4341127-4381724 (GRC m39). Each exon is represented by the same colour in different isoforms. Sizes of the exons are to scale. Top two lines show the representation of all exons present at this locus according to their order in the genomic sequence as e1, e2 etc. Line 2 indicates the nucleotide positions of each exon in a scenario where all the exons present. The exons have been arranged in linear order. (*B*) RT-PCR amplification of *Pirmy* showed multiple amplicons in both testis and brain, with many more amplicons in testis compared to brain. The RT-PCR products from brain and testis were cloned and sequenced to identify the splice variants. NTC is the non-template control. (*C*) BLAST analysis against mouse genome localizes the splice variants of *Pirmy* to NT_166343.2.

**Figure 3.**
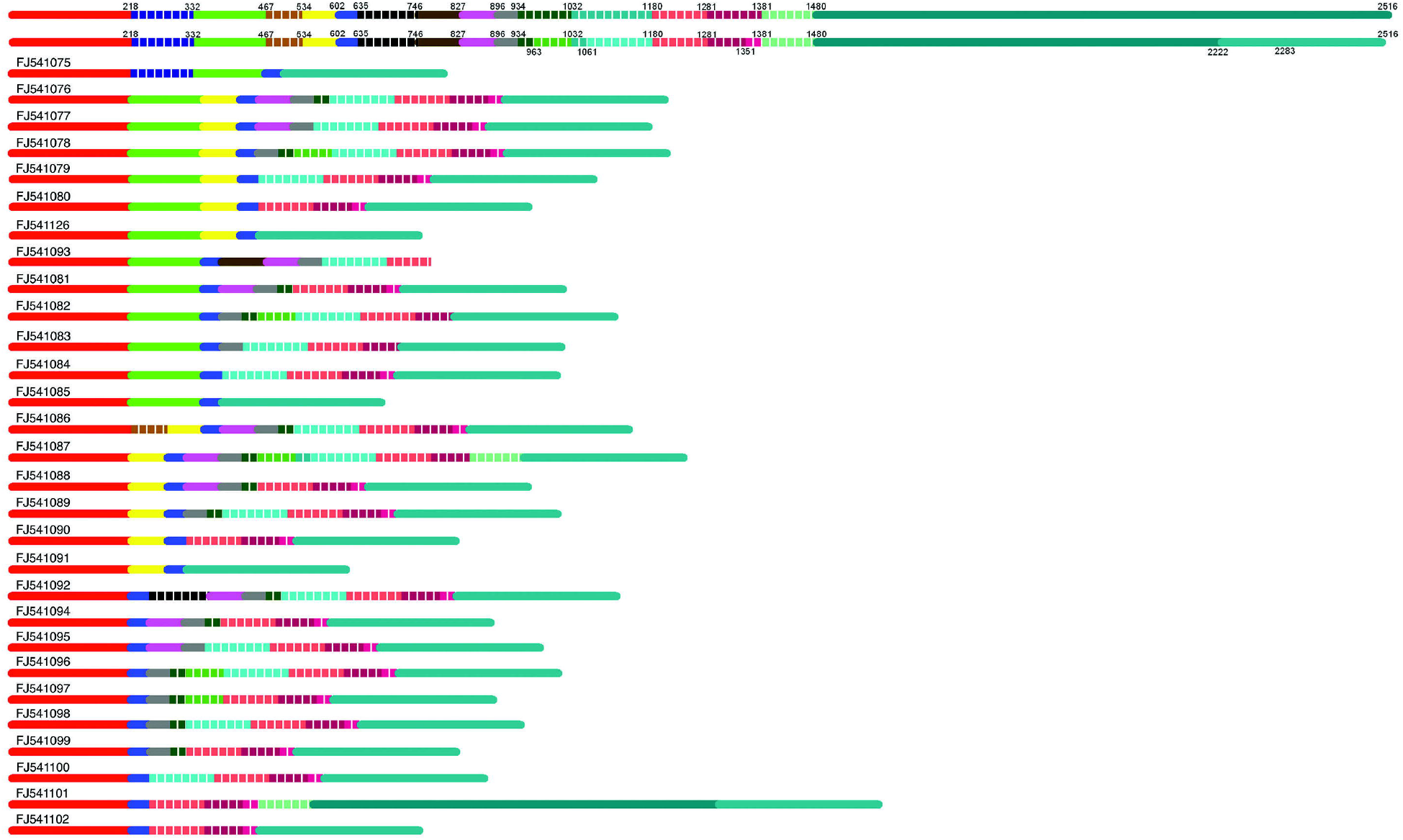
Localization of 28 transcript variants. (*A*) Top two lines show the representation of all exons present together according to their order in the transcript variants as e1, e2 etc. Putative nucleotide positions in a scenario wherein all the exons are present are indicated. Different splice variants have been arranged in linear order. Exons in dashed lines are specific to these 28 transcript variants, whereas other exons are common to *Pirmy* splice variants (Figure 2) and the transcript variants.

### Splice variants of *Pirmy* in mouse brain

Many transcripts that are expressed in testis are also known to be expressed in brain; therefore, we studied the expression of *Pirmy* in mouse brain also. RT-PCR amplification (Fig. 2*B*), cloning and sequencing of *Pirmy* products from mouse brain yielded 12 transcripts. All these transcripts localize to the single locus, to which *Pirmy* and its splice variants localized (4341127-4381724 (GRCm39)). In fact, comparison of the exons of the splice variants from the brain and testis showed that the same splice isoforms were present in both the tissues (Fig. 4).

**Figure 4.**
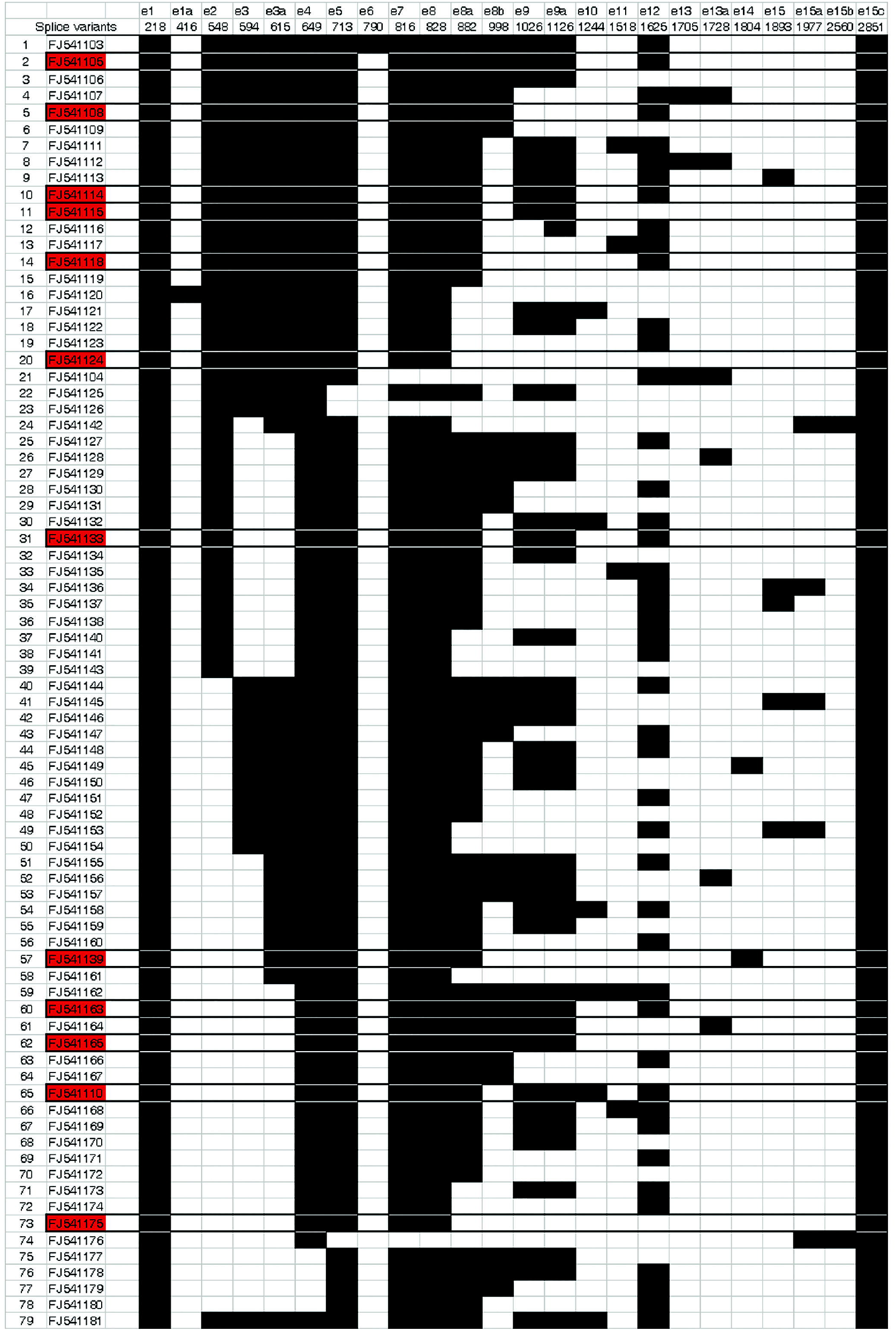
Comparison of splice-variants from testis and brain. The sequential exons from *Pirmy* splice-variants in testis have been represented in an Excel data sheet. Highlighted in red are the splice variants that are transcribed from male brain, which were also found in testis. The representation of e1, e2 etc. and the nucleotide positions is same as in Figure 2A.

### Expression of M34 in XY^RIII^qdel mice

Metaphase spreads from the XY^RIII^qdel mice showed a reduction in copy number of M34 on the Y chromosome showing that it localizes to the deleted region of the Y chromosome in the above mice (Fig. S3*B*). Therefore, expression of M34 was then checked in testis and sperms of XY^RIII^ and XY^RIII^qdel mice by FISH. Dramatic reduction in fluorescence intensity was observed in testis and sperm of XY^RIII^qdel mice. However, sperms from epididymis of both XY^RIII^ and XY^RIII^qdel mice showed faint fluorescence intensity (Fig. S3*B*). The subclones of M34 (Fig. S3A) when used as probes on testis sections showed reduction in fluorescence intensity in XY^RIII^qdel mice (Fig. 3*C*). We also checked the copy number of *Pirmy* in genomic DNA isolated from the wild type and XY^RIII^qdel mice by Real-Time PCR using primers to exon 7; a significant reduction in copy number was observed in the XY^RIII^qdel genome (Fig. S4C).

### Many proteins coded by autosomal genes are deregulated in XY^RIII^ qdel sperm proteome

We then analysed the motility profile of sperms from XY^RIII^ and XY^RIII^qdel mice and identified a stark difference in motility patterns (Movies S1, S2 respectively). Spermatozoa from XY^RIII^ mice show linear progressive motion whereas sperms from XY^RIII^qdel mice show rapid flagellar movement with non-linear and non-progressive motion. Most of the spermatozoa from XY^RIII^qdel mice stall at the same position with no linear displacement. Studies from Burgoyne’s lab reported morphological abnormalities in sperms from XY^RIII^qdel mice (Conway et al., 1994). In order to understand the connection between the Y-deletion and sperm abnormalities, we performed comparative sperm proteome analysis between normal mice and the XY^RIII^qdel mice by 2D-PAGE and mass spectrometry using protocols standardized in the laboratory (Bhattacharya, Devi, Dhople, & Jesudasan, 2013).

This analysis identified 8 protein spots that were differentially expressed in the pI ranges of 4-7 and 5-8 (Fig. 5*A*). Surprisingly four of these i.e. calreticulin (D1), Cu/Zn superoxide dismutase (SOD (D4)), fatty acid binding protein 9 (FABP9 (D5)), Serine Peptidase Inhibitor (Kazal type II (SPINK2)/Acrosin-Trypsin inhibitor variant 2 (D2) were upregulated in XY^RIII^qdel sperms compared to XY^RIII^ sperms (Fig. 5*A*). A novel shorter isoform of SPINK2 - SPINK2 variant3 (D3, Q8BMY) which was shorter by 27 amino acids at the N-terminal end was downregulated in XY^RIII^qdel sperms (Fig. 5*A*, Fig. S4B). Three proteins were not detectable in XY^RIII^qdel sperms in the pI range of 5-8 (Fig. 5A). Two of these were reported as hypothetical proteins in the NCBI database (1700001L19 Riken cDNA and 1700026L06 Riken cDNA. As we have identified these proteins, they have now been deposited in the Uniprot database with accession numbers Q9DAR0 and, Q7TPM5 (MAST (Bhattacharya et al., 2013) respectively. The third one was Stromal cell derived factor 2 like 1 (SDF2L1 (B)). Expression of four of the eight differentially expressed proteins was also confirmed by western blotting in testis and sperms (Fig. 5C).

**Figure 5.**
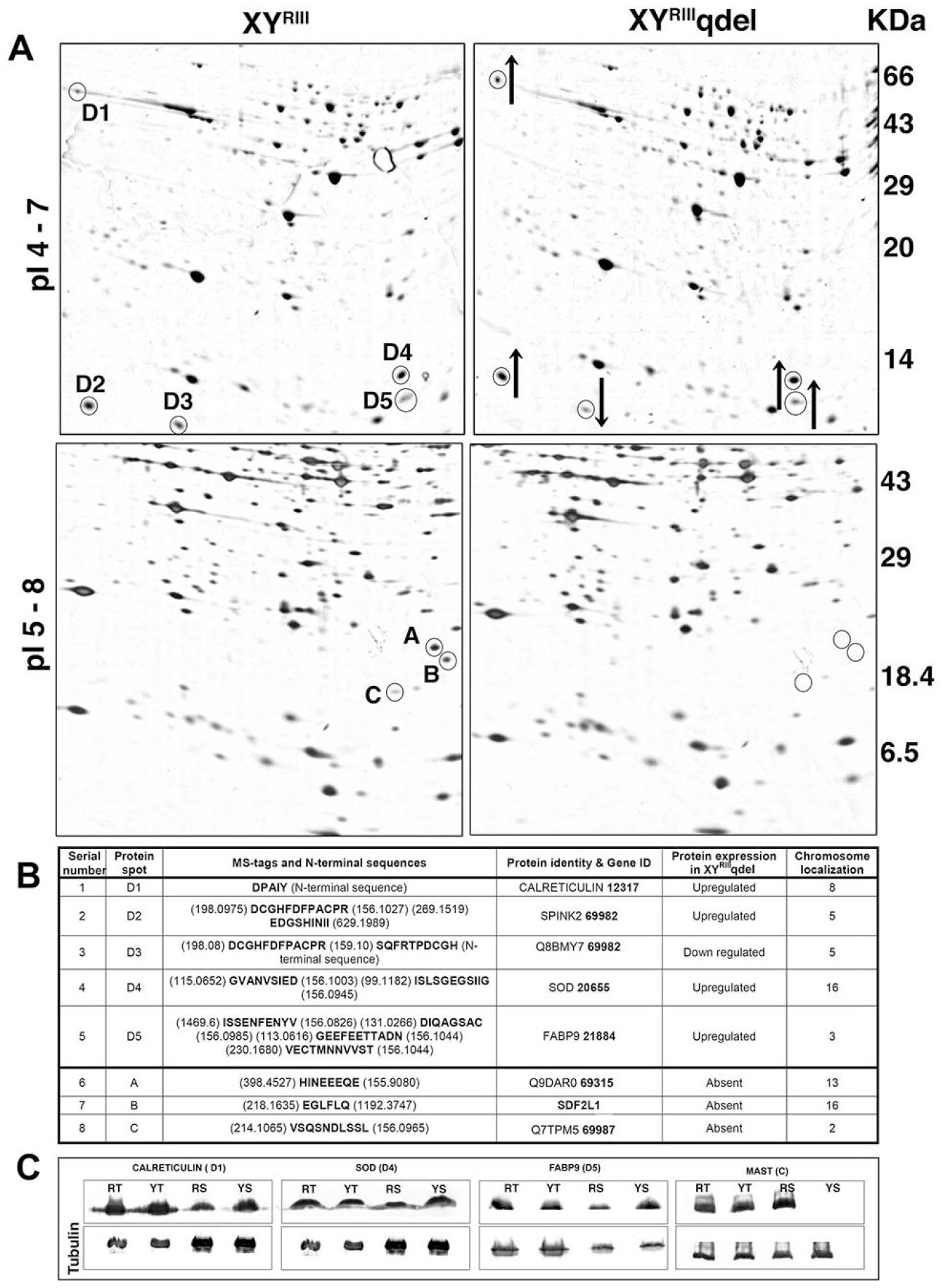
Sperm proteins are deregulated in XY^RIII^qdel mice. (*A*) Sperm lysates from the wild type XY^RIII^ strain and the mutant XY^RIII^qdel mice were separated by 2D-PAGE in the pI ranges of 4-7 and 5-8 on 8-20% gradient gels. Five differentially expressed proteins (D1 to D5) were identified by Mass Spectrometry analysis in the 4-7 pI range, of which 4 were upregulated (upward arrow) and one downregulated (downward arrow). Three proteins (A, B & C) were not detectable in the 5-8 pI range in XY^RIII^qdel compared to the XY^RIII^ sperm lysate. (*B*) The list of all the differentially expressed proteins identified in the proteomics screen is given in the table. The genes corresponding to all the differentially expressed proteins in XY^RIII^qdel localized to autosomes. (*C*) confirms the expression levels of four proteins (D1, D4, D5 and C) identified on 2D-gels by western blotting; the differential expression is observed in sperms but not in testis (RT - XY^RIII^ testis, YT - XY^RIII^qdel testis, RS - XY^RIII^ sperm, YS - XY^RIII^qdel sperm). The lower sub-panel for all blots is the loading control using Tubulin.

Calreticulin, SOD and FABP9 showed upregulation in XY^RIII^qdel sperms by both 2D PAGE and western blot analyses; MAST was not detectable by both the techniques in sperms. Protein expression of calreticulin, SOD, FABP9 and MAST did not vary significantly between testes of XY^RIII^ and XY^RIII^qdel mice. Thus, in sperms from the XY^RIII^qdel, four proteins were upregulated while one was downregulated. Three proteins were not visible in the XY^RIII^qdel sperm proteome. Surprisingly, all the eight genes corresponding to the differentially expressed sperm proteins localized to different autosomes (Fig. 5B).

Next, we analysed the expression of the transcripts corresponding to the protein spots in testis by Real-Time PCR/Northern blot analysis (Fig. S4A, B). Although the proteins Q9DAR0, SDF2L1 and MAST were not detectable in sperms of X^YRIII^qdel, the corresponding RNAs were present in testis. The transcripts of SDF2L1, MAST, calreticulin and SPINK2 variant 2 proteins were upregulated in XY^RIII^qdel mice testis. In contrast transcripts of Q9DAR0 and Spink2 variant 3 did not show significant quantitative difference between the two (Fig. S4B).

### UTRs of deregulated autosomal genes show homology to 108 transcripts

The fact that a few autosomal genes were deregulated when there was a deletion on the Y chromosome prompted us to investigate the mechanism behind this puzzling observation. We hypothesized that *Pirmy* transcript variants that show reduction in copy number in the genome and reduced transcription in XY^RIII^qdel, could be regulating autosomal gene expression in testis. BLAST analysis of these transcripts against the 3’ and 5’ UTRs of the deregulated genes revealed short stretches of homology ranging in size from 10-16 nucleotides, in either +/+ or +/- orientations. BLAST analysis of the 108 transcripts against the UTRs of deregulated genes identified 21 different hits. Of these 11 were from *Pirmy* splice variants and 10 from the other transcript variants respectively (Fig 6). There are as many as 7 *Pirmy* hits in the 3’ UTR of Q9DAR0 (Fig. 6). Homology between Y-derived transcripts and UTRs of deregulated autosomal genes indicates interactions between genes on the Y chromosome and autosomes in mouse testis. A BLAST against the entire transcripts of the deregulated genes showed homology to the coding regions also; however, more stringent BLAST parameters, showed homology to the UTRs alone.

**Figure 6.**
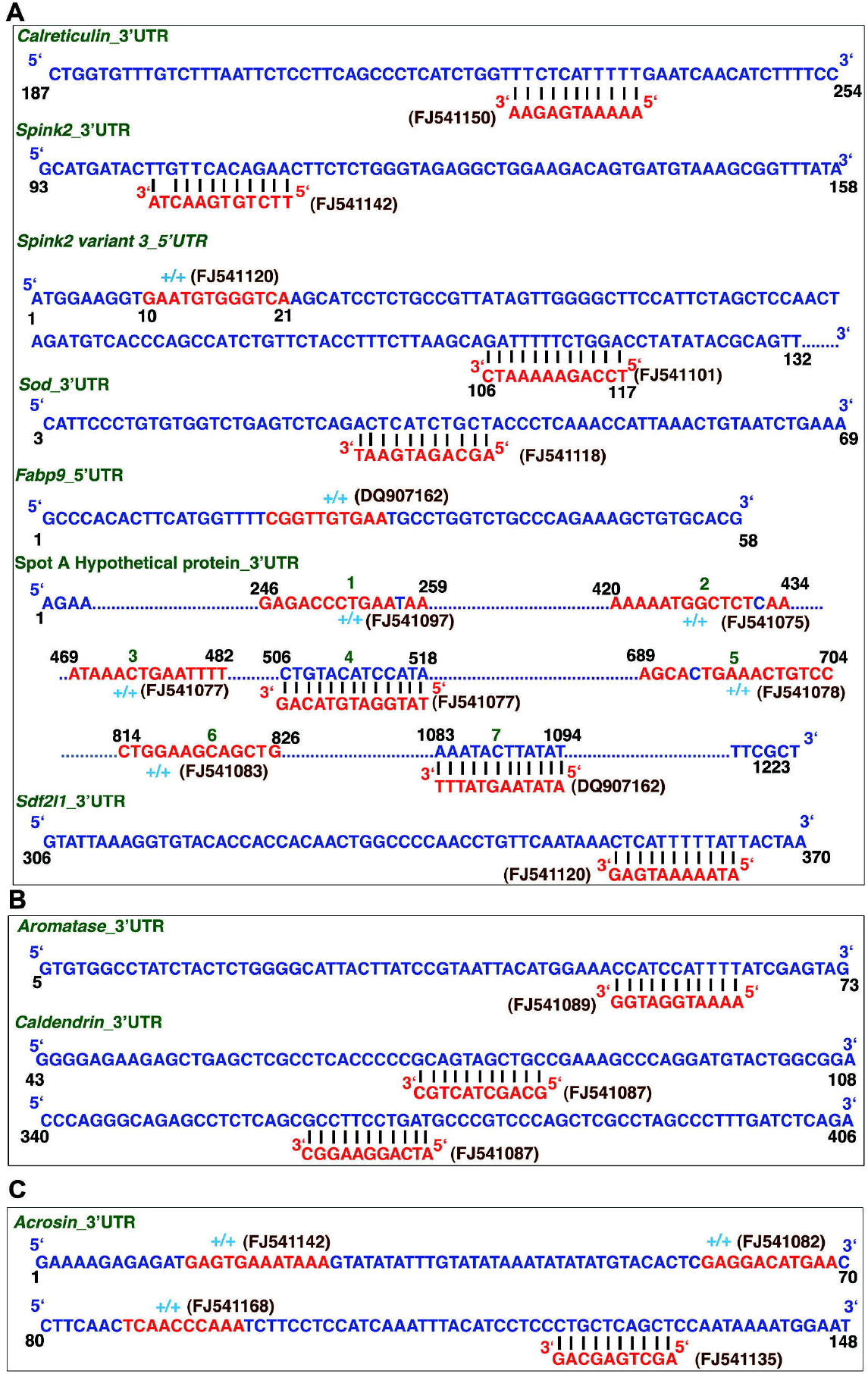
Localization of *Pirmy* homologous sequences to UTRs of deregulated genes. Panel (*A*) shows the UTR regions of the deregulated genes with the sequences homologous to *Pirmy* highlighted in red. Both +/+ and +/- homologies are observed. The splice isoforms of *Pirmy* are indicated in brown and the gene names in green color. (*B*) Couple of transcripts (aromatase and caldendrin) were identified independent of the proteomics screen (*C*) acrosin (literature survey) also harbor homologies to *Pirmy*.

Furthermore, we performed BLAST analysis of all the 108 transcripts (ncRNAs) against the entire UTR database. This identified small stretches of homology in the UTRs of a number of genes across different species. The homologous sequences (10-22 nt) localized to both exons and exon-exon junctions of the 108 transcripts. Representation of some UTR homologies are shown in Fig. 7A. We identified 372 unique homologous stretches in the 108 transcripts. Of these 302 (81.19%) localized to the exons and 70 (18.82%) to exon-exon junctions (Fig. 7B). The autosomal genes bearing homology to these ncRNAs in their UTRs, are expressed in multiple tissues including testis/epididymis (Table S1).

**Figure 7.**
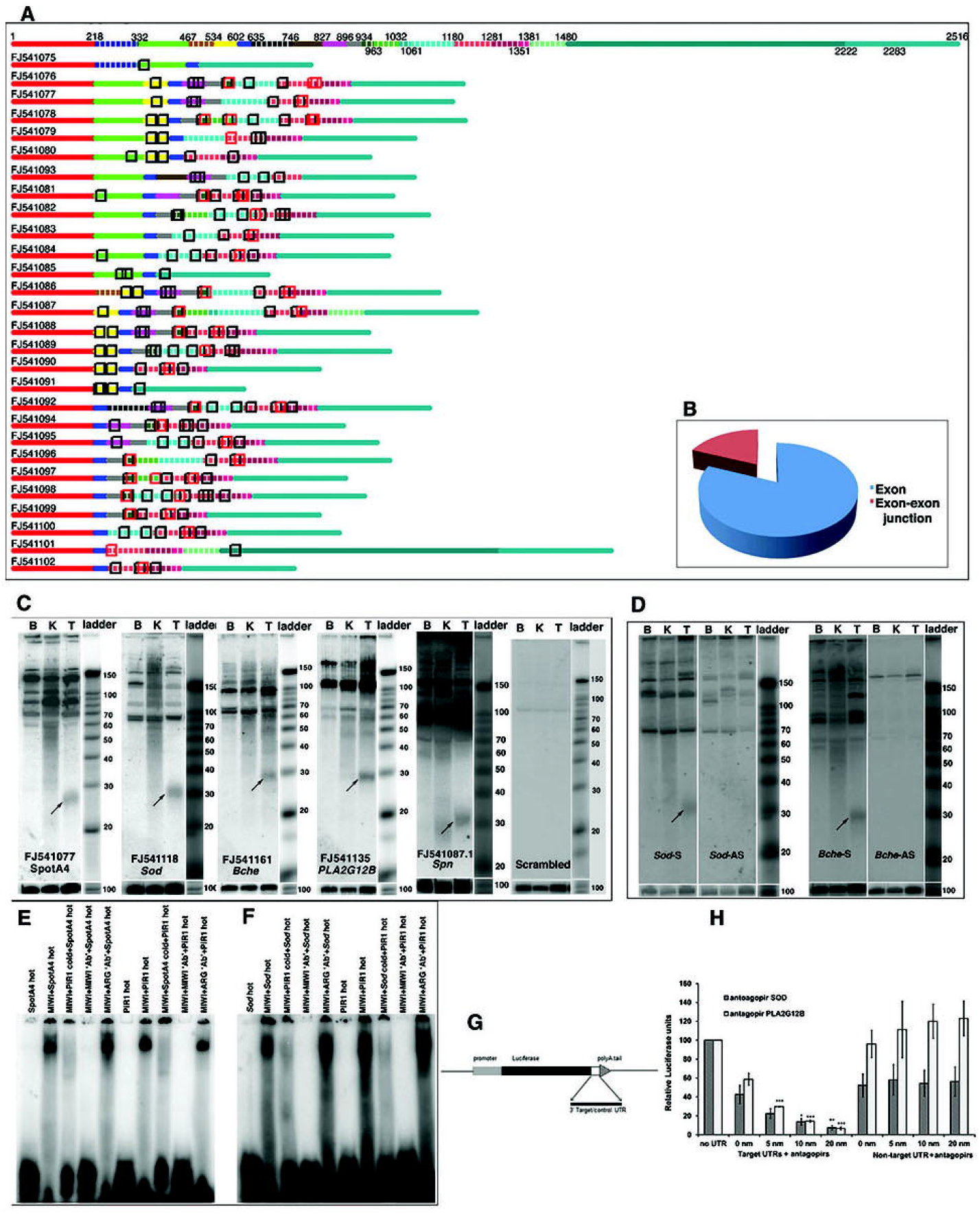
Identification of piRNAs in the transcript variants. (*A*) Short stretches of homology identified in the UTRs by BLAST analysis against UTR database are represented on the set of 28 transcript variants. The boxes highlighted in black match within exons and boxes highlighted in red match at the exon-exon junctions. Line 1 indicates the nucleotide positions as in Fig. 3*A*. (*B*) Pie chart shows ∼81% of the matches in exons and ∼19% at exon-exon junctions. (*C*) Use of sequences homologous to the 3’ UTRs of a hypothetical protein (SpotA4), *Sod, Bche, PLA2G12B* and *Spn* as probes on small RNA northern blots shows testis-specific signals of ∼30 nt size (indicated by arrows), which correspond to the size of piRNAs. Control blot using a scrambled oligonucleotide as probe shows no signal at ∼30 nt size. (*D*) Hybridization using probes from the sense (S) and antisense (AS) strands of *Sod* and *Bche* shows differential transcription from the two strands under the same conditions. Lower panels (C, D) show loading control using U6 probe. (*E, F*) EMSA using chemically synthesized RNA oligonucleotide sequences from *FJ541077* (E) and *FJ541118* (F) that have homology to the UTRs of the genes of hypothetical protein spot A (A4) and *Sod* respectively. These oligonucleotides and piR1 (Girard et al. 2006), a known piRNA, showed the shift in mobility with recombinant MIWI protein. The gel-shifts obtained with RNA-oligonucleotides from FJ541077 (spotA4) and FJ541118 (*Sod*) were competed out by cold piR1 and vice versa, i.e., the gelshift observed with piR1 was competed out by cold RNA-oligonucleotides from FJ541077 and FJ541118. Pre-incubation with MIWI antibody abolished the gelshift whereas pre-incubation with argonaute 3 antibody (ARG) did not - indicating specificity of binding. (*G*) Schematic representation of 3’UTR reporter constructs. (*H*) Results of reporter assay: Antagopirs to piRNA homologous sequences present in 3’UTRs of two genes, *Sod* and *PLA2G12B*, bring about concentration dependent reduction in luciferase expression. Use of these oligos against a non-target 3’UTR (*Cdc2l1*) did not affect luciferase expression.

### Identification of ∼30nt RNAs from *Pirmy* transcript variants

To check if these short stretches of homologies corresponded to small RNAs, few representative oligonucleotide sequences from the ncRNAs with homology to different UTRs were used as probes (Fig. S5) on small RNA northern blots. Two genes were chosen from the deregulated proteins Q9DAR0 (Spot A) and superoxide dismutase (SOD); Four genes (butyrylcholinesterase (*Bche*), phospholipase A2, group XIIB *(PLA2G12B) and* sialophorin *(Spn)*)were chosen from the BLAST output against UTR database. All the above probes elicited approximately 26-30 nt long testis-specific signals, of the size of piRNAs (Fig. 7*C*).

To confirm that these homologous sequences are indeed piRNAs, different experiments were designed. As the antiparallel strands of DNA are reported to express different levels of piRNA, differential expression from the antiparallel strands were studied using Sense (S) and antisense (AS) probes designed to homologous stretches in the 3’ UTRs of *Sod* and *Bche*. Identical experimental conditions showed differential expression of these 30 nt species of RNAs from the two strands of DNA (Fig. 7*D*), further indicating that these short RNAs could be piRNAs.

As piRNAs are PIWI/MIWI binding small RNAs, Electrophoretic Mobility Shift Assay (EMSA) using the *Pirmy*-derived oligonucleotides and recombinant MIWI protein was done to check if the sequences with homologies to UTRs of different genes are indeed piRNAs. Representative gel shifts using oligonucleotides from UTRs of Q9DAR0 and *Sod* are depicted in Fig. 7E and F respectively. piR1 (Girard, Sachidanandam, Hannon, & Carmell, 2006), a known piRNA served as the positive control. Specificity of binding was indicated by the use of corresponding cold competitors as described in the legend to Fig. 7E and F. The piRNA derived oligonucleotides competed out binding of piR1 to MIWI protein and vice versa. This confirmed that these oligonucleotides are indeed MIWI-binding RNAs and therefore piRNAs. The mobility shift was also competed out by MIWI antibody while Argonaute 3 antibody did not alter the mobility of the gel-shifted band obtained with MIWI indicating specificity of binding. These experiments provide further evidence that these short RNA sequences are piRNAs.

### *Pirmy* identifies piRNAs from NCBI Archives database

Next line of evidence to the fact these short stretches of homologies are piRNAs came from the NCBI Archives database for piRNAs. BLAST analysis using the 108 transcripts identified a total of 1445 small RNAs in the Sequence Read Archives SRP001701 and SRP000623 with >95% identity. These transcripts identified 352 MIWI2 - MILI-associated reads in SRP000623. Of these 310 had homology to midrepetitive sequences (SINEs, LINEs and LTRs) and 5 had homology to X chromosome. Thus, the remaining 37 piRNAs could be specifically derived from the Y chromosome.

### Antagopirs downregulate reporter gene expression

Complementary oligonucleotides synthesized to piRNA sequences present in UTRs of *Sod* and *PLA2G12B* were designated as antagopirs (Fig. S5). Cloning of these UTRs 3’ to the Luciferase gene reduced its expression. Treatment with increasing concentrations of antagopirs, i. e. 5 nM, 10 nM and 20 nM caused further concentration dependent reduction in Luciferase expression (Fig. 7*H*). The antagopirs to *Sod* and *PLA2G12B* did not have an effect when the UTR from a non-target gene (*Cdc2l1)* was cloned 3’ to the Luciferase gene. Fig. 7G is a schematic representation of 3’UTR reporter construct. Thus, the use of antagopirs to piRNAs could modulate gene expression in vitro.

This study therefore, paved the way for a series of novel and exciting observations. We have identified, a novel, polyadenylated lncRNA (*Pirmy*) expressed from mouse Yq in testis, that shows the large number of splice variants in testis. These Y-derived transcripts harbour piRNAs that have homology to the UTRs of a few autosomal genes expressed in mouse testis. The proteins expressed from these autosomal genes are deregulated in sperms of Y-deleted mice and appear to be regulated by piRNAs generated from MSYq derived lncRNAs. Consolidation of the results from the proteomics analysis and the molecular studies suggested that piRNAs generated from male-specific mouse Yq regulate expression of multiple autosomal genes in testis. Partial deletion of Yq resulted in deregulation of these proteins leading to sperm anomalies and subfertility. Thus, subfertility in mice appears to be a polygenic phenomenon that is regulated epistatically by the Y chromosome.

## DISCUSSION

Y-chromosomes harbour genes for male determination and male fertility. Yet the role of Y chromosomal repeats in male fertility and spermatogenesis remains enigmatic. In this study, we elucidate the role in male fertility of a species-specific repeat, M34, from mouse Y long arm, which is transcribed in mouse testis. Transcription from repeats on mouse Y chromosome has been reported earlier. Testis-specific transcription of a family of poly(A)RNAs from the mouse Y chromosome was first reported by Bishop and Hatat using a multicopy Y-derived probe, pY353/B (Bishop & Hatat, 1987; Conway et al., 1994). Subsequently more transcripts were identified in mouse testis, using repeat sequences localizing to mouse Y chromosome (Prado et al., 1992; Toure et al., 2005). The report of *Pirmy* transcript variants described in this study adds to the repertoire.

The 108 *Pirmy* transcript variants in the present study were discovered serendipitously by cloning and sequencing of the multiple RT-PCR products obtained using primers to the initial and terminal exons. As the primers were restricted to just two of the exons, it is possible that we might discover more transcript variants using primers to different combinations of exons for RT-PCR amplification. Alternative splicing has been reported in ncRNAs (Pang et al., 2005), yet, such extensive splicing as observed in our study has not been reported for any of them. Very few polymorphically spliced genes have been described earlier from sex chromosomes, particularly the Y (Ellis, Ferguson, Clemente, & Affara, 2007; Wegmann, Dupanloup, & Excoffier, 2008). Thus, the 80 splice variants of *Pirmy* in this study appears to be by far the maximum number of isoforms characterized from a single ncRNA by alternative splicing. Elucidation of identical splice isoforms in mouse brain and testis in this study (Figs. 2, 4) shows that these alternative splicing events are precise and not random because the same splicing events take place in both the tissues. The consensus splice signal sequences present at the intron-exon junctions in all the splice variants further reaffirm programmed splicing events.

The identification of deregulated proteins, for which the corresponding genes localise to autosomes, in a Y-deletion mutant was a surprise. Identification of small stretches of homology in the UTRs of these transcripts from the *Pirmy* transcript variants established the connection between the two. Observation of 27-32 nt long signals on small RNA northern blots using the short homologous sequences as probes, which also bind to MIWI protein and show differential expression from the antiparallel strands of DNA indicated that these probes identify piRNAs. Some of these also identified piRNAs in the SRA databases. Small RNAs of the size of ∼27-32 nucleotides that bind PIWI protein are classified as piRNAs. These are also known to be differentially transcribed from the two strands of DNA. The presence of piRNAs in the UTRs of genes corresponding to deregulated proteins, suggests the regulation of these autosomal genes by Y-derived piRNAs. The concentration dependent reduction of Luciferase expression by antagopirs corroborates the regulation of these genes by Y-derived piRNAs. Proteins from three other autosomal genes are also deregulated in different strains of Y-deleted mice besides the ones identified in the proteomics screen; caldendrin is upregulated in XY^RIII^qdel sperm (Bhattacharya et al., 2013) and acrosin is downregulated in B10.BR-Y^del^ sperm. (J. Styrna, Klag, & Moriwaki, 1991). Aromatase is overexpressed in both the Y-deleted strains of mice i.e. XY^RIII^qdel (our unpublished observation), and B10BR-Y^del^ (Kotula-Balak, Grzmil, Styrna, & Bilinska, 2004). Therefore, it is not surprising to find *Pirmy* homologous sequences in the UTRs of caldendrin, acrosin and aromatase genes, suggesting Y-mediated regulation for these genes as well. Ellis and colleagues also observed up or down regulation of genes from the X-chromosome and autosomes in testes of mice with deletions of Y heterochromatin using a microarray approach (Ellis et al., 2005). Homology between UTRs of some of the above genes and the 108 ncRNAs (Table S2) further strengthens the hypothesis of putative regulation of genes located elsewhere in the genome by Y chromosomal repeats.

piRNAs are known to regulate gene expression at the levels of both transcription and translation. Our study also suggests regulation at both the transcriptional and post-transcriptional levels by the piRNAs derived from the 108 transcript variants. Genes corresponding to three proteins that were missing from the mutant sperm proteome, were indeed transcribed in testis. The fact that there is comparable expression of calreticulin and SPINK2 variant2 proteins in XY^RIII^ and XY^RIII^qdel testes by western blot analysis, despite significantly upregulated expression of these transcripts in testis, could indicate regulation at the level of both transcription and translation for these genes. piRNAs regulate translation in early embryos and gonads besides containing transposable elements (Aravin, Hannon, & Brennecke, 2007; Deng & Lin, 2002).

We also propose that the deregulated proteins identified in the current study are at least partially responsible for the sperm phenotype observed in XY^RIII^qdel mice. For example, FABP9 that is upregulated in XY^RIII^qdel spermatozoa, localizes to perforatorium, the subacrosomal region in spermatozoa with falciform head shapes (Korley, Pouresmaeili, & Oko, 1997; Oko & Morales, 1994; Pouresmaeili, Morales, & Oko, 1997). It is the most abundant protein of the perforatorium, and putatively has a role in shaping the unique sperm head structure of rodents (Oko & Morales, 1994; Pouresmaeili et al., 1997). The deregulated expression of Spink2 isoforms also could contribute to sperm head morphological abnormalities and reduced sperm motility; Kherraf and colleagues (Kherraf et al., 2017) observed grossly misshapen sperm heads and reduced motility in Spink2 knock out mice.

Calreticulin overexpressed in XY^RIII^qdel spermatozoa is a calcium store associated with sperm functions such as hyperactivated motility, capacitation and acrosome reaction (Yanagimachi, 1982). The subsequent cascade of events could result in subfertility. Caldendrin yet another protein that is upregulated in XY^RIII^qdel sperm (Bhattacharya et al., 2013), localizes to acrosome in rats and is considered to be a stimulus dependent regulator of calcium (Redecker, Kreutz, Bockmann, Gundelfinger, & Boeckers, 2003). SPINK2 variant3 localizes to the acrosome in mouse spermatozoa (our unpublished results). The physiological function of SPINK2 is believed to be blocking of deleterious degradation of proteins released by acrosin from spermatozoa during acrosome reaction (Moritz, Lilja, & Fink, 1991). Being a putative sperm acrosin inhibitor a role in fertility can be envisaged. *Sdf2l1* is an ER stress-inducible gene, induced by the unfolded protein response pathway (Fukuda et al., 2001). The XY^RIII^qdel spermatozoa may be more susceptible to stress induced damages as they lack stress response protein, SDF2L1.

MAST a novel protein that is not visible in the XY^RIII^qdel sperm proteome, but localizes to both the acrosome and sperm tail, indicating a role in sperm motility and penetration of the egg (Bhattacharya et al., 2013). Bioinformatic analysis of the novel protein Q9DAR0 predicts it as a cilia related gene (McClintock, Glasser, Bose, & Bergman, 2008). Based on this information putative role of this protein in sperm motility can be envisaged. SOD, a protein upregulated in XY^RIII^qdel, has been found to be positively associated with sperm count and overall motility (Lu, Huang, & Lu, 2010). The irreversible conversion of androgens to estrogens is catalyzed by aromatase transcribed by *Cyp19* gene (Simpson et al., 1994). Biology behind the skewed sex ratio towards females in the progeny sired by XY^RIII^qdel males could be explained by the upregulated expression of aromatase in testes of these mice. Acrosin plays a crucial role in acrosome exocytosis and egg zona pellucida penetration (Yamagata et al., 1998). Spermatozoa lacking acrosin (acrosin^-/-^) exhibit delayed fertilization as both the processes of acrosome exocytosis and egg penetration are deferred (Adham, Nayernia, & Engel, 1997; Mao & Yang, 2013).

Functions of Y chromosome have been elucidated using different deletions of the chromosome in the past. Naturally occurring deletions in the euchromatic long arm of Y chromosome in azoospermic men showed the involvement of this region in human male infertility (Vogt, 1998). *Drosophila melanogaster* males with deletions of different regions of the Y chromosome show absence of several sperm axoneme proteins (Goldstein, Hardy, & Lindsley, 1982). Mice with partial or total deletions of Y heterochromatin show deregulation of testicular gene expression and subfertility/sterility (Cocquet et al., 2009; Ellis et al., 2005). Previous studies in the lab elucidated another example of an intronless Y-derived ncRNA mediated regulation of an autosomal gene, *CDC2L2* via trans-splicing in human testis (Jehan et al., 2007). The role of mouse Y heterochromatin in the current study therefore reveals a novel pathway for the regulation of autosomal genes by Y chromosome, mediated by piRNAs, in male reproduction. Therefore, consolidation of the observations from mouse and human studies therefore shows that Y chromosome regulates autosomal genes expressed in testis using distinct mechanisms in different species.

Comparative sperm proteomics analysis in our study portrays involvement of multiple autosomal genes in subfertility. The regulation of autosomal gene expression appears to be relaxed in sperms of Yq-deleted mice. This reflects a connection between the Y chromosome and autosomes. In fact, as suggested by Piergentili, Y chromosome could be a major modulator of gene expression (Piergentili, 2010). Our results seem to provide explanation for some of the earlier classical observations of mice with different Y chromosomal deletions exhibiting subfertility/sterility along with sperm morphological abnormalities, fewer motile sperms, sex ratio skewed toward females etc. Similar phenotypes are also observed in cross-species male sterile hybrids of *Drosophila* and mouse (Albrechtova et al., 2012; Campbell, Good, Dean, Tucker, & Nachman, 2012; Heikkinen & Lumme, 1998; Johnson, Hollocher, Noonburg, & Wu, 1993; Piergentili, 2010; Tao, Zeng, Li, Hartl, & Laurie, 2003; Vigneault & Zouros, 1986; White, Stubbings, Dumont, & Payseur, 2012). Y-chromosome has also been implicated in the male sterility phenotype of these interspecies hybrids (Campbell et al., 2012; Carvalho, Vaz, & Klaczko, 1997; Lamnissou, Loukas, & Zouros, 1996; Vigneault & Zouros, 1986; Zouros, Lofdahl, & Martin, 1988). Thus, the phenotypes observed in cross species male sterile hybrids and the Y-deletion mutants are comparable. Introduction of Y-chromosomes into different genetic backgrounds of *Drosophila* resulted in deregulated expression of hundreds of genes localizing to the X-chromosome and autosomes (David et al., 2005; Lemos, Araripe, & Hartl, 2008). It has also been proposed that incompatibility between the Y-chromosomes and different autosomes could result in the hybrid dysgenesis of sperm related phenotypes observed in *Drosophila* (Zouros et al., 1988). Zouros and colleagues also suggested the presence of epistatic networks in interspecies hybrids, based on the fact that homospecific combination of alleles at a given set of loci could sustain normal development, but heterospecific combinations could not (Davis, Noonburg, & Wu, 1994; Lamnissou et al., 1996; Wu & Palopoli, 1994). This early hypothesis seems to be amply supported by our study. Further, our results elucidate the Y - derived piRNAs as the genetic basis of epistatic interactions between Y chromosome and autosomes in mouse. Our results also suggest for the first time, the mechanism of piRNA mediated regulation of autosomal genes involved in spermiogenesis and male fertility. This is, to our knowledge is the first report on possible regulation of autosomal genes involved in male fertility and spermiogenesis, mediated by Y-encoded small RNAs/piRNAs in any species.

In brief, the XY^RIII^qdel mutant strain of mouse, where there is a partial deletion of long arm of the Y chromosome, exhibit sperm morphological and motility related aberrations and subfertility (Conway et al., 1994). A comparative sperm proteomic profiling of the XY^RIII^ and XY^RIII^qdel mice captured few differentially expressed proteins that could partially account for the aberrant sperm phenotype. Surprisingly, genes corresponding to the deregulated proteins localized to autosomes and not to the deleted region of the Y chromosome. Earlier we demonstrated an event of trans-splicing between a Y-ncRNA and a protein coding autosomal mRNA in human testis for putative translational regulation. A search for the Y-autosome connections in mouse led to the identification of novel ncRNAs from mouse Y long arm that subsequently was shown to regulate the genes expressed in testis via piRNAs. Thus, adopting a top-down approach we have established a novel mode of regulation of autosomal genes expressed in mouse testis by the Y chromosome and the biology behind the aberrant sperm phenotype in Y-deleted mice.

Finally, evolutionary impact of novel genetic interactions or regulatory mechanisms such as those reported in this study could be significant. The generation of piRNAs from species-specific repeats on mouse Y-chromosome that apparently regulate autosomal gene expression in testis raises more questions in the field of speciation and evolution. Do mutations in the Y chromosomal repeats collapse the poise of the species? Are species-specific repeats on the Y chromosome the fulcrum on which rests the fine balance between species identity and evolution?

## METHODS

### Animals and Reagents

The XY^RIII^ strain (wild type) and the XY^RIII^qdel strain (Y-deletion mutant) of mice used in the study was a gift from Prof. Paul S Burgoyne, MRC, UK. All the animals used in the experiments were bred and reared in the in-house animal facility of our institute (CCMB), in accordance with the guidelines from Indian Science Academy under CPCSEA (Committee for the Purpose of Control and Supervision of Experimental Animals). This study was approved by the Institutional Animal Ethics Committee (IAEC 65/28/2006). The recombinant construct in pAAV-IRES-hrGFP from which MIWI protein was isolated, was a gift from Arvind Kumar, CCMB, Hyderabad.

### Identification of cDNA using M34 (DQ907163)

An amplified mouse testis cDNA library (Mouse testis MATCHMAKER cDNA Library – Clonetech) was screened with the male specific ES cell EST (CA533654) with sequence homology to the genomic clone DQ907163 (Fig. 1*E*) as per manufacturer’s protocol. Mouse testis cDNA Library was screened according to standard protocol at a stringency of 2 x SSC, 0.1% SDS at 65°C for 10 min. A total of 2 × 10^5^ colonies were screened to obtain 18 clones after tertiary screening. Male-specificity of the positive clones from the library was determined using Southern blots containing mouse male and female DNA. All 18 clones gave the same sequence. The sequence was submitted to the NCBI database and is named as *Pirmy* (DQ907162).

### Identification of Splice Variants

Total RNA (1 μg) isolated from brain and testes tissues each of XY^RIII^ strain of mouse were reverse transcribed with the (SuperscriptII, Reverse Transcriptase enzyme, Invitrogen), using oligo (dT) primers and random hexamers. The RT-mixes were concentration equalized using GAPDH primers. DQ907162 was amplified after two rounds of PCR using primers to the first and last exons. For the first round, forward (GTGTGACAGGGTGGGGAATC) and reverse primers (TTCCTGAAGATAGCACTTGTG), and the following conditions were used - initial denaturation 95°C, 1min, cycle denaturation 95°C, 1 min, annealing 62°C, 30 sec and extension 72°C for 2min (35cycles), final extension 72°C for 7 min. The second round of amplification was done using nested primers GAGGACCGTATTCATGGAAGAG (forward) and GCAAATGGCTCACATCAGTGG (reverse) using initial denaturation at 95°C for 1 min, cycle denaturation at 95°C, 1 min, annealing at 66°C for 30 sec and extension at 72°C for 2 min (38 cycles) and final extension at 72°C for 7 min. Annealing temperatures up to 70°C yielded multiple products. The multiple products obtained after two rounds of PCR from both testis and brain (Fig. 2*B*) were cloned into pCR TOPO vector using Invitrogen TOPO TA cloning kit. Approximately 1000 clones were sequenced on 3730 DNA Analyser (ABI Prism) using sequencing kit BigDye Terminator V3.1 Cycle Sequencing kit. These yielded 108 unique transcripts. BLAST analysis of these transcripts against the genomic sequences (GRCm39) at a stringency of >97% localised 80 of these to a single locus on the Y chromosome (4341127-4381724) and the rest to multiple sites on the Y.

Therefore, the 80 transcripts that localised to the single locus are splice variants of *Pirmy*.

### Sperm Proteome Analysis

Sperm lysates (1 mg of cell weight per 5 µl of lysis buffer – [Urea 8 M, CHAPS 4% (w/v), Tris 40 mM, Biolyte (3-10) 0.2% (w/v) and TBP (1µl/100 µl]). The suspension was incubated on ice for one hour to allow buffer to permeabilise and lyse the sample. Further, the sample was briefly sonicated on ice. The lysate was centrifuged for 15 min at 13,000 rpm at 4°C. The supernatant was collected and was further taken for ultra-centrifugation at 55,000 rpm for one hour at 4°C. The clear lysate was collected in fresh tube and PMSF added to a final concentration of 1mM. The protein concentration in the cell lysate was estimated by Bicinchoninic acid assay (Pierce, Rockford IL), following manufacturer’s instructions in a micro titre plate. BSA was used as the standard for estimation. The proteins were separated in the first dimension on 4-7 and 5-8 IPG strips (Bio-Rad). The strips were then loaded onto 8-20% gradient PAGE for separation on the second dimension. Protein spots were visualized by Coomassie Blue staining. Spot to spot matches were done to identify differences. Analysis of five sets of gels after normalization with control spots using PDQUEST software version 6.0 (Bio-Rad) identified the differential proteins. Measuring the optical density of these differentially expressed proteins in arbitrary units validated the quantitative differences (Fig. S7). These values were subjected to nonparametric Kruskal-Wallis H test and the levels of confidence determined by the Chi-squared test (75-95% degrees of confidence). Trypsin digested spots were processed to obtain the protein tags by MS analysis on Hybrid Quadrupole TOF mass spectrometer (API QSTAR PULSAR i, PE SCIEX).

### Small RNA Isolation

Total RNA was extracted from male kidney, testis and brain of mice using TRIZOL reagent (Invitrogen). Total RNA was denatured at 65°C for 10 min, incubated with 10% PEG-8000 (Sigma-Aldrich) and 5 M NaCl for 30 min on ice and centrifuged at 7,000 rpm for 7-10 min. The supernatant containing small RNA was precipitated overnight with 3 volumes of absolute alcohol and centrifuged at 13,000 rpm for 30 min. The small RNA pellet was washed with 80% ethanol and resuspended in RNase free Gibco water. The quality of small RNA was checked on 12% Urea PAGE (Biorad) and quantitated using Nanodrop V-1000 (Thermo Fisher Scientific).

### Small RNA Northern Blotting

20-50 µg of small RNA from each tissue was resolved on a 12% Urea PAGE gel in 0.5 x TBE running buffer and transferred onto Hybond N^+^ membrane. Decade marker (Ambion) was labelled and loaded according to the manufacturer’s instructions. 10-25 µM of each LNA-oligonucleotide probe (Exiqon), was end labelled for use as probes (hybridization buffer - 5 x SSC, 5 x Denhardt’s and 1% SDS). Blots were hybridized at 37°C and washed from 37°C to 65°C in 2 x SSC, 0.2% SDS depending on the intensity of the signal. U6 was used as the loading control. Fig. S6 shows the location of the LNA (locked nucleic acid) probes used for small RNA northern blots on the corresponding splice variants.

### Real-time PCR analysis

#### Copy number estimation of *Pirmy*

Genomic DNA was isolated from testes of XY^RIII^ and XY^RIII^qdel mice using phenol-chloroform method and quantified using a Nanodrop (NANODROP 2000, ThermoScientific). Quantitative Real-time PCR (LightCycler 480, Roche) was performed using SYBR green master mix (Qiagen) with 2 ng of genomic DNA and a primer concentration of 200 nM per reaction. The primers used were as follows: *Pirmy* (exon 7): forward 5’-GTG CGG TTG TGA AGG TGT TC– 3’, reverse 5’-CCT CCA CCT TCC ATT CAC CC-3’; *Gapdh:* forward 5’-ACG GGA AGC TCA CTG GCA TGG −3’, reverse 5’-CAA CAG CGA CAC CCA CTC CTC–3’. PCR conditions included an initial denaturation for 5 min at 95°C followed by 45 cycles of denaturation at 95°C for 10 secs, annealing at 58°C for 20 sec and elongation at 72°C for 30 sec. The amplification of specific product was confirmed by melting curve profile (cooling the sample to 65°C for 1 min and heating slowly with an increase in temperature of 5°C at each step till 95°C, with continuous measurement of fluorescence). The relative fold change in *Pirmy* was analyzed based on Livak method (2^-ΔΔCt^).

### Real-time PCR analysis of differentially expressed genes

The total RNA was extracted from testes using Trizol (Invitrogen). One μg of RNA was reverse transcribed to cDNA using Verso cDNA synthesis kit (ThermoScientific). qPCR was performed using SYBR green master mix (Qiagen) and analysed in Roche Light Cycler LC480. The primer sequences corresponding to mouse cDNA used were as follows: forward 5’-CGA GGG CCA GAC AGG GAT TG–3’ and reverse 5’-CCC ATA GAC AGA GGA CAT CAG-3’ for Riken cDNA 1700001L19; forward 5’-ACT TCC CGT CGC CGC TAT C-3’ and reverse 5’-TGA CCG ACA GGA ACA CAG AGG-3’ for *Sdf2l1*; forward 5’-CAG CAT CGA GCA GAA GTA TAA GC-3’ and reverse 5’-TGG GTG GAG TTA TTG CAG TAG-3’ for *Mast*; forward 5’-GGA AAC CAC GTC AAA TTG −3’ and reverse 5’-GGT GAT GAG GAA ATT GTC-3’ for Calreticulin; forward 5’-GGC TAC TTG ACC ACT GC-3’ and reverse 5’-TTT GAG AAT CGG AAG AGT C-3’ for *Spink2* Variant 2; forward 5’-TTC CGA ACA CCA GAC TG-3’ and reverse 5’-ATG GCT ACC GTC CTC C-3’ for *Spink2* Variant 3; forward 5’-TGA AGT CGC AGG AGA CAA CCT-3’ and reverse 5’-ATG GCC TTC CGT GTT CCT A-3’ for *Gapdh*. PCR conditions included an initial denaturation for 5 min at 95°C followed by 45 cycles of denaturation at 95°C for 10 secs, annealing at 58°C for 20sec and elongation at 72°C for 30 sec. The amplification of specific product was confirmed by melting curve profile. The relative fold change in expression was estimated based on Livak method (2^-ΔΔCt^).

### Electrophoretic mobility shift assay

RNA oligonucleotides corresponding to GAAGCAGAUGAGUAUAUG from *Sod* and UCAUUGGACAUAAACUGAAUUUUCCA from the gene for hypothetical protein spot A (Q9DAR0) were end labelled with γ-^32^P ATP and column purified using G-25 Sephadex (Sigma-Aldrich) and quantitated on a scintillation counter. EMSA reactions were set up in a total volume of 25 μl using binding buffer (20 mM HEPES, 3 mM MgCl2, 40 mM KCl, 5% Glycerol, 2 mM DTT and RNase inhibitor (4U)), with MIWI protein (5 μg per reaction). MIWI was over-expressed from a recombinant construct in pAAV-IRES-hrGFP vector and purified using the FLAG tag. Competitors i.e., unlabeled oligonucelotides (30 x concentration of hot oligo), MIWI antibody (90 ng) and Argonaute 3 antibody (100 ng) were added to the reaction, incubated for 1 h on ice, before addition of radio-labeled oligonucleotide (7000-10,000 cpm) and the entire mix was incubated on ice for another 30 min. EMSA was done on 5% native PAGE and image captured using FUJI phosphor Imager (FUJIFILM FLA-3000). A known piRNA, piR1, was used as the positive control, Argonaute 3 antibody served as the antibody control.

### Luciferase assay

Either Luciferase gene alone or luciferase along with the UTR was cloned into pcDNA3.1 expression vector for assaying the effect of antagopirs (Figure 7H) on Luciferase expression. Co-transfection experiments were done using the GC-1spg cell line (ATCC CRL-2053) and lipofectamine 2000 (Invitrogen) using protocols specified by the manufacturer. Cells were seeded in 48-well plates, 24 hrs prior to transfection to obtain approximately 80% confluency. Each well was transfected with 50 ng of pcDNA3.1 plasmid containing either Luciferase gene alone or along with the cloned UTRs, 50 ng of the β-gal plasmid and varying concentrations of oligonucleotides complementary to the piRNA along with 0.5µl of lipofectamine 2000 in antibiotic and serum free DMEM (GIBCO). The complementary oligonucleotide to the piRNA has been designated as antagopirs. The antagopirs (*Sod* - 5’GAAGCAGAUGAGUAUAUG3’; *PLA2G12B* - 5’CCAAACUGUUGGAAGAAGGAAU3’) were procured as RNA oligonucleotides from Eurofins Genomics India Pvt. Ltd, Bangalore, India. Different concentrations of antagopirs (0 nM, 10 nM, 20 nM and 40 nM/ well) were tested in the assay for their effect on the UTRs. Five hrs post transfection, the medium was replaced with complete growth medium. The cell extracts were prepared 24 hrs post transfection using Reporter Lysis Buffer (Promega) and assayed for Luciferase activity in EnSpire 2300 multimode plate reader (Perkin Elmer). The Luciferase activity was normalized using β galactosidase. Three independent sets of experiments were done in triplicates.

## DATA ACCESS

All sequences from this study have been deposited in NCBI database. With the accession numbers: DQ907162.1, FJ541075-FJ541181, Q7TPM5, Q9DAR0, Q8BMY7.

## ACKNOWLEDGEMENTS

We would like to dedicate this manuscript to Prof. Lalji Singh and Prof. Burgoyne both of whom have left us for their heavenly abode and whom we miss greatly at this point of time. We gratefully acknowledge the gift of the XY^RIII^ and XY^RIII^qdel mice by Prof. Paul S Burgoyne, MRC, Edinburgh, UK. This study could not have been done without the gift of these mice. The GC-1spg cell line was gifted by Prof. MRS Rao, JNCASR, Bangalore, India. We thank Dr. Dinesh Kumar and Professor B. K. Thelma for reading the manuscript and giving useful inputs and Mr. Sivarajan Karunanithi for help with partial *in silico* analysis.

The funding by Department of Science and Technology, India (SP/SO/B70/2001) and Department of Biotechnology, India (BT/PR 10707/AGR/36/596/2008), intramural funding from Council of Scientific and Industrial Research (CSIR), India to RAJ and fellowships by CSIR, India to HMR, RB are acknowledged.

## Author contributions

HMR, KM did experiments and partial bioinformatics analysis. RB, ZJ, PA, NMP, VMD, MS, BS, JLA, SMT, RRD, SMR performed experiments. ST, AC, SK contributed to *in silico* analysis. NR helped with confocal imaging. LS gave the probe and inputs. RAJ conceived, guided the work and wrote the manuscript.

